# BCAS3 regulates WIPI2b via a direct interaction in the ATG16L1-binding cleft

**DOI:** 10.64898/2026.11.06.764939

**Authors:** Philippa Rosewell, Alina Üffing, Laura Masino, Sharon A. Tooze

## Abstract

*De novo* formation of the phagophore membrane is a fundamental step in autophagy and relies on recruitment of the E3-like ligase complex via an interaction between ATG16L1 and phosphatidylinositol 3-phosphate (PtdIns3P)-effector protein, WIPI2b. Although WIPI2b can directly bind membranes, how its membrane association is regulated is still not fully understood. Here, we describe BCAS3 as a novel direct interactor of WIPI2b and demonstrate that a peptide of BCAS3 binds in the ATG16L1-binding cleft of WIPI2b with low micromolar affinity. We show that BCAS3 and its partner PHAF1 induce recruitment of WIPI2b to membranes via the interaction of BCAS3 with PtdIns3P. This not only challenges the current paradigm, whereby ATG16L1 is the only confirmed binder of WIPI2b in this region, but also offers a mechanism through which WIPI2b membrane association might be regulated.

## Introduction

Canonical bulk/non-selective macroautophagy (hereafter autophagy) is dependent on sequential activation and recruitment of protein complexes which help to build a *de novo* double-membraned organelle termed the phagophore that sequesters cargo to be subsequently degraded in the lysosome [1]. A key signal within this process is the formation of phosphatidylinositol 3-phosphate (PtdIns3P) by the PtdIns3-kinase complex I (PtdIns3K; composed of VPS34, p150, Beclin 1 and ATG14 [2,3]) which results in the recruitment of PtdIns3P-effector proteins including DFCP1 [4] and the WIPI proteins [5]. There are four mammalian WIPIs (WIPI1-4) which are included in the family of proteins called PROPPINS; PROPPINS are characterised by beta-propeller folds, PtdIns binding motifs, and an amphipathic helix which can insert into a membrane [6–10]. WIPI1 and WIPI2 are the mammalian orthologues of yeast Atg18 while WIPI3 and WIPI4 are more closely related to Hsv2 [11]. The WIPIs have distinct but not fully understood roles. WIPI4 preferentially interacts with the lipid transfer protein ATG2 [12] and is thought to be important for later stages in autophagy where it regulates autophagosome size [13,14]. WIPI2, on the other hand, is nearly essential for autophagy [14] and functions mainly in early phagophore formation where it recruits the E3-ligase complex (ATG16L1-ATG12-ATG5) thereby promoting lipidation of ATG8 family members [11,15–17]. It has become evident that localization of WIPI2 to the phagophore is not simply regulated by the formation of PtdIns3P but also by post-translation modifications [9] and recruitment by protein-protein interactions in selective autophagy [18,19]. Crucially, WIPI2 membrane association seems unaffected by starvation [11] and there is still residual membrane association in a WIPI2b mutant lacking its amphipathic helix [9] suggesting that there are additional factors regulating WIPI2 recruitment and function.

We previously discovered ATG16L1 and the rest of the E3-like complex (ATG12 and ATG5) in the interactome of WIPI2b [15]. In addition to these key autophagy proteins, several other interactors were identified including Breast Carcinoma Amplified Sequence 3 (BCAS3) and c16orf70 (also referred to as Mytho or Phagosome Assembly Factor 1 [PHAF1]) which, at the time, had no annotated function in autophagy. Since discovering BCAS3 and PHAF1 in the interactome of WIPI2b, the literature linking these proteins to autophagy has increased [20–24], although there has been little consensus on the mechanism underlying their role.

The functions of these proteins are unclear although both appear to have important roles in health and disease. PHAF1 is a FoxO-dependent gene which has been shown to regulate skeletal muscle health in mouse models [23] and contribute to healthy ageing in *C. elegans* [22]. Similarly, PHAF1 KO zebrafish show alterations to lifespan and motor function [25]. Crucially, although PHAF1 KO models show alterations to autophagy, and it has been proposed to act as a scaffold with WIPI2 [22], it is not clear how this is related to the phenotypes described. BCAS3 was originally identified as a gene amplified in breast cancer [26] and early work described it as cytoskeleton-related protein important for angiogenesis and embryogenesis [27–29] with Bcas3 KO mice showing embryonic lethality at E9.5 [28]. Loss of BCAS3 in zebrafish results in neurodevelopmental defects [30], a phenotype which is recapitulated in human patients with BCAS3 loss of function mutations; affected patients show global developmental delay, microcephaly and short stature [31]. More recently, BCAS3 has been described to associate with the phagophore alongside PHAF1 [20,21,24] and WIPI1/WIPI2/ATG-18 [21,24] although how it functions in autophagy has not yet been elucidated. Previous work has generally focussed on non-selective autophagy although there may be some selectivity of function; BCAS3 KO had no effect on valinomycin-induced mitophagy in mammalian cells [21], however selective autophagy of paternal mitochondria in *C. elegans* embryogenesis relied on BCAS-3 and PHAF-1 [24]. Structurally, BCAS3 also resembles a PROPPIN, containing a WD40 domain and PtdIns3P binding motif [20], which may relate it functionally to the WIPI proteins. Interestingly, magnetic resonance imaging of patients with BCAS3 mutations showed features resembling brain iron accumulation [32]; this is a key feature in patients with beta-propeller associated neurodegeneration (BPAN), a disease caused by mutations to the WIPI4 gene, *WDR45*, and related to WIPI4’s autophagic function [14,33,34].

Given the emerging literature linking BCAS3 and PHAF1 to WIPI2 function, and autophagy more generally, we set out to characterise the mechanism through which these proteins act. Here, we present BCAS3 as a novel direct interactor of WIPI2b which is required for effective recruitment of WIPI2b to the nascent phagophore membrane and an important regulator of autophagy.

## Results and Discussion

### BCAS3 recruits WIPI2 to the phagophore

BCAS3, PHAF1 and WIPI(1/2) have been shown to co-immunoprecipitate with one another [15,20–22] and the interaction between BCAS3 and PHAF1 is conserved across species including *Dictyostelium discoideum* [20], *Caenorhabditis elegans* [22], and *Homo sapiens* [21,22] suggesting an important function of this complex. We wanted to observe the colocalization of this protein complex in cells and so used the “split Venus” bimolecular fluorescence complementation assay [35,36] to allow visualisation of the three components at once. We generated fusion proteins where the Venus N-terminal domain (V1) was fused to the N-terminus of PHAF1, and the Venus C-terminal domain (V2) was fused to the N-terminus of BCAS3, while non-fused V1 (V1-ORF) served as a control. If the fused proteins of interest interact, the V1 and V2 domains reform the complete Venus fluorophore and emitted fluorescence can be detected. HEK293A WT cells were transfected with V1-PHAF1 and V2-BCAS3, and autophagy was induced with 2 hours of culture in Earle’s Balanced Salt Solution (EBSS). Punctate Venus signal could be observed indicating an interaction between PHAF1 and BCAS3 (**Figure 1A**). We observed only diffuse cytosolic Venus signal when V1-ORF and V2-BCAS3 were expressed confirming the specificity of the Venus signal (**Fig. S1A**). Immunofluorescence with an antibody against WIPI2 showed co-localization of WIPI2 and Venus (PHAF1+BCAS3) confirming that WIPI2, BCAS3, and PHAF1 interact during starvation (**Figure 1A**). These regions of colocalization were mostly punctate, but we also observed complex ring-like structures which were positive for all components.

**Figure 1.**
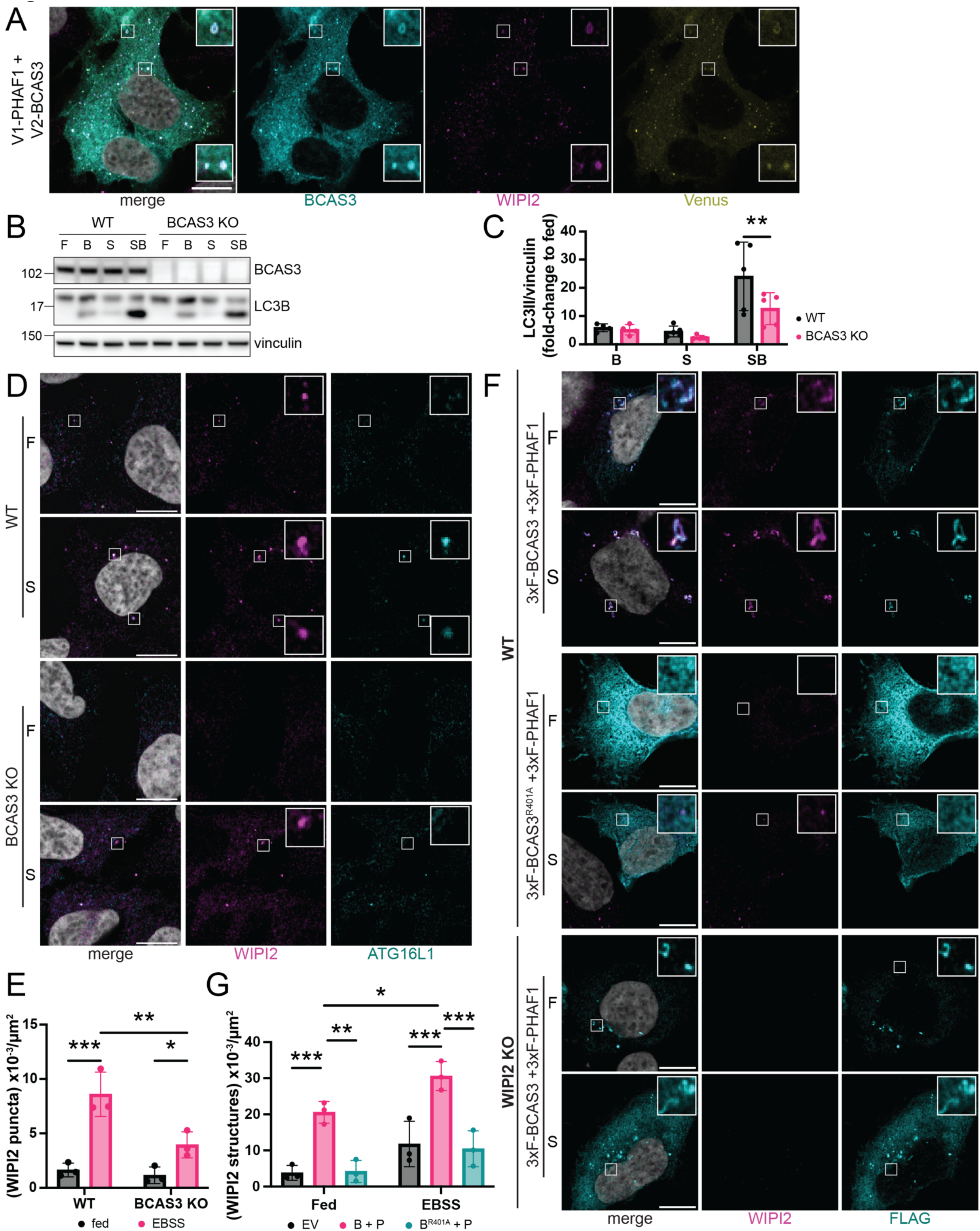
BCAS3 recruits WIPI2 to the phagophore. (**A**) Representative images of HEK293A WT cells transiently transfected with V1-PHAF1 and V2-BCAS3 (Venus, yellow). Cells were treated with EBSS for 2 hours then stained with antibodies against BCAS3 (cyan) and WIPI2 (magenta). Example regions of co-localisation are shown as inset zooms. Scale bar, 10 µm. (**B**) Representative western blot of HEK293A WT or BCAS3 KO cells cultured in complete media (F) or EBSS (S) for 2 hours with and without 100 nM Bafilomycin A1 (B and SB, respectively). (**C**) Quantification of (B). n = 4-5 as indicated, two-way ANOVA followed by Fisher’s LSD post-hoc test. **p = 0.0052. (**D**). Representative images of HEK293A WT or BCAS3 KO cells cultured in complete media (F) or EBSS (S) for 2 hours and then stained with antibodies against WIPI2 (magenta) and ATG16L1 (cyan). Regions of interest are shown as inset zooms. Scale bar, 10 µm. (**E**) Quantification of WIPI2 puncta density per area of cell cytoplasm from (D). n = 3, two-way ANOVA followed by Fisher’s LSD post-hoc test. *p = 0.0279, **p = 0.0022, ***p = 0.0002. (**F**) Representative images of HEK293A WT or WIPI2 KO cells transiently transfected with 3xFLAG-PHAF1 (3xF-PHAF1) and either 3xFLAG-BCAS3 WT (3xF-BCAS3) or 3xFLAG-BCAS3^R401A^ (3xF-BCAS3^R401A^). Cells were cultured in complete media (F) or EBSS (S) for 2 hours and then stained with antibodies against WIPI2 (magenta) and FLAG (cyan). Regions of interest are shown as zooms. Scale bar, 10 µm. (**G**) Quantification of density of WIPI2-positive structures as seen in (F). HEK293A WT cells were transiently transfected with 3xFLAG empty vector control (EV), 3xFLAG-BCAS3 WT + 3xFLAG-PHAF1 (B + P) or 3xFLAG-BCAS3^DRER^ + 3xFLAG-PHAF1 (B^R401A^ + P). Cells were cultured in complete media (Fed) or EBSS for 2 hours and stained as in (F). n = 3, two-way ANOVA followed by Tukey’s post-hoc test. *p = 0.0118, **p = 0.0011, EV fed vs B + P fed ***p = 0.0009, EV EBSS vs B + P EBSS ***p = 0.0003, B + P EBSS vs B^R401A^ + P EBSS ***p = 0.0002.

BCAS3 contains a WD40 domain and has been proposed to be a new member of the PROPPIN family [20], of which the other mammalian family members are WIPI1-4. Given the importance of WIPIs in autophagy, we postulated that BCAS3 may be a key interactor of WIPI2 and therefore generated HEK293A BCAS3 KO cells using CRISPR/Cas9 gene editing to further examine this relationship. Successful editing was validated by western blot as well as amplicon sequencing (**Fig. S1B**). We assessed autophagy flux by measuring lipidation of ATG8 paralogue, LC3B (to form LC3B-II). Cells were treated for 2 hours in either complete media (F) or subjected to serum starvation with EBSS (S). To inhibit lysosomal degradation of LC3B-II, cells were also treated with Bafilomycin A1 (B and SB). BCAS3 KO cells showed a significant reduction in accumulation of LC3B-II following EBSS and Bafilomycin A1 compared to WT, indicating that loss of BCAS3 impairs autophagy (**Figure 1B and 1C**). We further examined autophagic processing by utilising the Halo-LC3 processing assay [37] which specifically shows degradation of LC3 in the lysosome and provides a more downstream readout. Autophagy was induced in stable HEK293A WT^Halo-LC3B^ or BCAS3 KO^Halo-LC3B^ with incremental time points of starvation (0, 2, 4, or 6 hours). In WT^Halo-LC3B^ cells, free Halo could be observed with a maximum at 4 hours (**Fig. S1C**), while in BCAS3 KO^Halo-LC3B^ cells processing after 4 hours EBSS treatment showed a significant reduction in (**Fig. S1D**) further supporting a role for BCAS3 in normal autophagic function.

It has previously been described that loss of PHAF1 reduces formation of WIPI2 puncta [22]; given that PHAF1 and BCAS3 colocalize with WIPI2b as well as each other (**Figure 1A**), we wondered whether the impairment we observed in LC3B lipidation and processing was due to an effect on WIPI2 function. Cells were treated with EBSS for 2 hours to induce autophagy, and immunofluorescence was used to examine formation of early autophagic structures using antibodies against WIPI2 and ATG16L1 (**Figure 1D**). In WT cells, starvation results in a significant increase in the density of WIPI2 puncta (**Figure 1D and 1E**). In BCAS3 KO cells EBSS treatment still resulted in a small increase in WIPI2 puncta formation, but the puncta density was significantly reduced compared to WT (**Figure 1D and 1E**). This was also accompanied by a reduction in ATG16L1 puncta, due to a loss of WIPI2-mediated recruitment.

Finally, we wanted to examine the functional relationship between BCAS3 and WIPI2. Since loss of BCAS3 resulted in a reduction of WIPI2 puncta, we reasoned that overexpression of the BCAS3-PHAF1 complex should increase WIPI2 puncta formation. HEK293A WT cells were transiently transfected with 3xFLAG-BCAS3 and 3xFLAG-PHAF1 (3xF-BCAS3 + 3xF-PHAF1) or an empty vector control (3xF-EV) and examined by immunofluorescence (**Figure 1F and Fig. S1E**). We observed that overexpression of the BCAS3-PHAF1 complex increased formation of WIPI2 clusters following starvation, and in fact was sufficient to cause a significant increase in WIPI2-positive structures under fed conditions (**Figure 1F and 1G**). These structures included rings, cups and extended lines which likely represent membranous structures. BCAS3 has previously been shown to contain PtdIns3P-binding motifs suggesting it may directly bind the phagophore membrane [20,21]. We therefore expressed a 3xFLAG-tagged PtdIns3P-binding mutant (3xF-BCAS3^R401A^ + 3xF-PHAF1) which was previously reported to prevent BCAS3 puncta formation [21]. We observed a significant reduction in FLAG-positive structures in both fed and starved conditions (**Fig. S1F**) but also observed that WIPI2 was no longer recruited into clusters in either fed or starved conditions (**Figure 1F and 1G**). We confirmed this was not due to differences in expression by western blot (**Fig. S1G**). Furthermore, some FLAG-positive clusters were positive for ULK1 (**Fig. S1H**) indicating that these represent autophagy related structures. During selective degradation of paternal mitochondria in *C. elegans*, the recruitment of BCAS-3 was impaired by loss of WIPI2 ortholog ATG-18 [24]. To test whether BCAS3 recruits WIPI2 or vice versa, the BCAS3-PHAF1 complex was overexpressed in HEK293A WIPI2 KO cells. FLAG-positive structures could still be observed under fed conditions and then significantly increased following 2 hours of serum starvation (**Figure 1F and Fig. S1F**); this is presumably due to direct recruitment to PtdIns3P-positive membranes as these structures were not observed with 3xF-BCAS3^R401A^ (**Fig. S1E**). We further confirmed that the interaction between BCAS3 and PHAF1 was independent of WIPI2b by using GFP-trap to immunoprecipitate GFP-BCAS3 from HEK293A WIPI2 KO cells starved for 2 hours (**Fig. S1I**). GFP-BCAS3 co-immunoprecipitated PHAF1 to a similar level in both WIPI2 KO and WT cells indicating that the interaction between BCAS3 and PHAF1 occurs upstream of their interaction with WIPI2b. We therefore propose a model whereby BCAS3 and PHAF1 form a stable complex, even under fed conditions, which then acts upstream of WIPI2b to enhance its recruitment it to PtdIns3P-rich membranes following induction of autophagy.

### BCAS3 binds WIPI2b in the ATG16L1 binding cleft

Given the evidence that BCAS3 and PHAF1 form a complex with one another, we wondered whether they might associate into a trimeric complex with WIPI2b. As BCAS3 and PHAF1 are relatively unstudied proteins there are currently no experimentally determined structures, so we used AlphaFold3 (AF3) [38] to examine whether BCAS3, PHAF1 and WIPI2b can form a trimeric complex (**Figure 2A**). A large interface was predicted between BCAS3 and PHAF1 which included contacts between a helical region covering amino acids 710-730 of BCAS3 and the C-terminus of PHAF1. This prediction complements published experimental data which showed that a deletion mutant of BCAS3 lacking amino acids 680-719 impairs the interaction between BCAS3 and PHAF1 [21] and therefore supports the validity of the AF3 prediction. We observed that, despite previous suggestions that PHAF1 may be a direct interactor of WIPI2 [22], there were no predicted interfaces between PHAF1 and WIPI2b, and an AF3 model containing only PHAF1 and WIPI2b had a very low ipTM score (0.2, data not shown). Instead, a C-terminal region of BCAS3 was predicted to form a helix which interfaces with WIPI2b. We therefore hypothesised that the BCAS3-PHAF1 complex recruits WIPI2b to the phagophore via a direct interaction between BCAS3 and WIPI2b.

**Figure 2.**
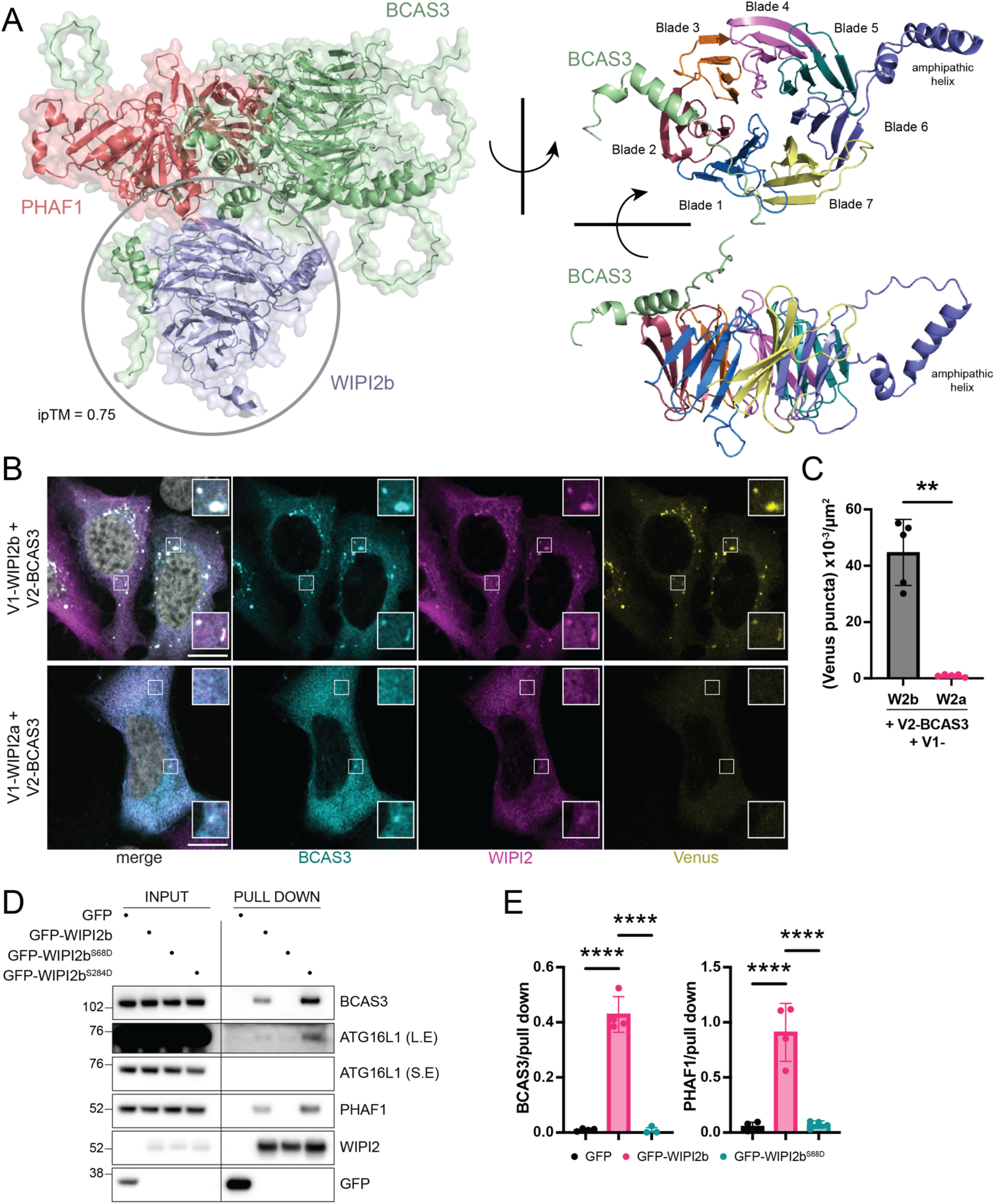
BCAS3 binds WIPI2b in the ATG16L1 binding cleft. (**A**) Left: AlphaFold3 model of PHAF1 (red), WIPI2b (purple) and BCAS3 (green). ipTM = 0.75. Right: zoom of circled region showing only WIPI2b beta-propeller (coloured by blade) and interacting region of BCAS3 (green) sitting in the cleft between blade 2 and blade 3. Visual representation based on [39] to highlight similarity to ATG16L1 binding mode. (**B**) Representative images of HEK293A WT cells transiently transfected with V2-BCAS3 and either V1-WIPI2a or V1-WIPI2b (Venus, yellow). Cells were treated with EBSS for 2 hours then stained with antibodies against BCAS3 (cyan) and WIPI2 (magenta). Regions of interest are shown as inset zooms. Scale bar, 10 µm. (**C**) Quantification of Venus puncta density per area of cell cytoplasm in transfected cells. n = 5, Welch’s t-test. **p = 0.0011. (**D**) Representative western blot of GFP-trap. HEK293A WIPI2 KO cells were transiently transfected with either a GFP empty vector control (GFP), GFP-WIPI2b WT (GFP-WIPI2b), GFP-WIPI2b^S68D^ or GFP-WIPI2b^S284D^ and cultured in EBSS for 2 hours. Input samples represent 1% of the total lysate. (**E**) Quantification of (D) with pull down of protein of interest normalised to WIPI2 pull down. n = 4, one-way ANOVA followed by Tukey’s post-hoc test. ****p < 0.0001.

We noted that in the AF3 model, the helix of BCAS3 was predicted to interact with WIPI2b between blade 2 and 3 of the beta-propeller; a region known to be important for the binding of ATG16L1, a key interactor of WIPI2 [15,39,40] (**Figure 2A**). Notably, there are multiple splice isoforms of WIPI2 (a-e) with WIPI2b generally being considered key for bulk autophagy [41]. WIPI2a shares complete sequence identity with WIPI2b except for a loop insertion which prevents interaction with ATG16L1 such that WIPI2a does not localize to the phagophore during starvation [15]. We therefore postulated that if BCAS3 interacted with WIPI2b in the ATG16L1-binding cleft, then BCAS3 would not interact with WIPI2a. We examined this using the split Venus assay where the V1 domain was fused to either WIPI2a or WIPI2b while the V2 domain was fused to BCAS3. HEK293A WT cells transfected with split Venus components were starved for 2 hours with EBSS before being examined by confocal microscopy. When V1-WIPI2b and V2-BCAS3 were expressed, we observed formation of Venus puncta (**Figure 2B**), which were also recognised by antibodies against BCAS3 and WIPI2, further confirming their interaction. There was a significant reduction in formation of Venus-positive puncta with expression of V1-WIPI2a and V2-BCAS3 (**Figure 2B and 2C**), despite detection of BCAS3 and WIPI2a expression with antibodies. Notably, formation of the complete Venus fluorophore is irreversible and therefore stabilises protein-protein interactions. Interestingly, stabilisation of the WIPI2b-BCAS3 interaction in this way resulted in formation of extended structures (i.e. non-punctate) in around 40% of transfected cells and highly elongated structures (**Fig. S2A**) in around 5% of transfected cells, suggesting that BCAS3 could positively regulate WIPI2b-mediated phagophore growth. To confirm that WIPI2a and BCAS3 cannot interact, HEK293A WIPI2 KO cells were transfected with a GFP empty vector, GFP-WIPI2a, or GFP-WIPI2b, and subjected to 2 hours of EBSS treatment followed by a GFP-trap to look at interactors of GFP-WIPI2b. As expected, ATG16L1, BCAS3, and PHAF1 were co-immunoprecipitated with GFP-WIPI2b while GFP-WIPI2a, on the other hand, showed no pulldown of ATG16L1, BCAS3 or PHAF1 (**Fig. S2B**). The loop insertion in WIPI2a therefore prevents interaction with the BCAS3-PHAF1 complex in line with the AF3 prediction that BCAS3 binds in the ATG16L1-binding cleft.

We have previously shown that phosphorylation of WIPI2b at S68 by the autophagy regulating kinase, ULK1, causes release of ATG16L1 from the WIPI2b binding pocket [9]. We reasoned that if the interaction between WIPI2b and BCAS3 occurs in this same pocket, then BCAS3 binding might also be regulated by this phosphorylation. HEK293A WIPI2 KO cells were transfected with a GFP empty vector, GFP-WIPI2b WT, GFP-WIPI2b^S68D^ or GFP-WIPI2b^S284D^ (another ULK1 phosphosite unrelated to ATG16L1 binding [9]). Autophagy was induced with 2 hours of EBSS treatment before a GFP-trap was performed with the various GFP-WIPI2b constructs. GFP-WIPI2b WT co-immunoprecipitated ATG16L1, BCAS3 and PHAF1 as expected, as did the GFP-WIPI2^S284D^ mutant (**Figure 2D**). Pulldown of GFP-WIPI2b^S68D^ showed a complete loss of co-immunoprecipitation of BCAS3 and PHAF1 (**Figure 2D and 2E**), in addition to the previously reported loss of ATG16L1 pulldown. This further supports the model that BCAS3 binds to the ATG16L1 binding cleft and suggests that phosphorylation by ULK1 could regulate the interaction between BCAS3 and WIPI2b.

It was previously proposed that the interaction between PHAF1 and WIPI2 was dependent on ATG16L1 as the interaction was lost when the WIPI2-RERE mutant (which is ATG16L1-binding deficient) was expressed [22]. To rule out that the interaction we were observing between BCAS3 and WIPI2 was not a result of ATG16L1 binding, we performed a GFP-trap in HEK293A ATG16L1 KO cells expressing either a GFP empty vector control or GFP-WIPI2b after 2 hours of EBSS treatment (**Fig. S2C**). GFP-WIPI2b was still able to co-immunoprecipitate BCAS3 and PHAF1 in the absence of ATG16L1. Our data therefore supports a model whereby BCAS3 interacts with WIPI2b in the ATG16L1 binding cleft, independently of ATG16L1.

### BCAS3 binds directly to WIPI2b with micromolar affinity

Having established that the BCAS3-WIPI2b interaction is dependent on the ATG16L1-binding cleft, we wanted to confirm that this was due to a direct interaction between BCAS3 and WIPI2b. This region has also been identified as a potential binding site for other WIPI2 interactors including the mitophagy receptor NIX [18,19] and core autophagy protein ATG13 [19] although no direct binding has been shown. We compared the binding interfaces of BCAS3-WIPI2b and ATG16L1-WIPI2b. Negatively charged residues on ATG16L1, E226 and E230, have been shown to be essential for its interaction with WIPI2 (via electrostatic interactions with K88 and R108 respectively) [15,39,40]. The crystal structure of a peptide of ATG16L1 bound to truncated WIPI2b [40] is recapitulated by using AF3 to model this region of ATG16L1 (aa 207-237) with WIPI2b (full length) (**Figure 3A**). We identified the corresponding region on BCAS3 and modelled a peptide of the same length (aa 845-875) which AF3 predicts to form a helix. This helix interacts with K88 and R108 of WIPI2b via corresponding negatively charged residues on BCAS3 (D864 and E868) (**Figure 3A**). AF3 also robustly predicts interactions between BCAS3 E855 and WIPI2b K128 (shown in sticks in **Figure 3A**) for both full-length BCAS3 and BCAS3^845-875^.

**Figure 3.**
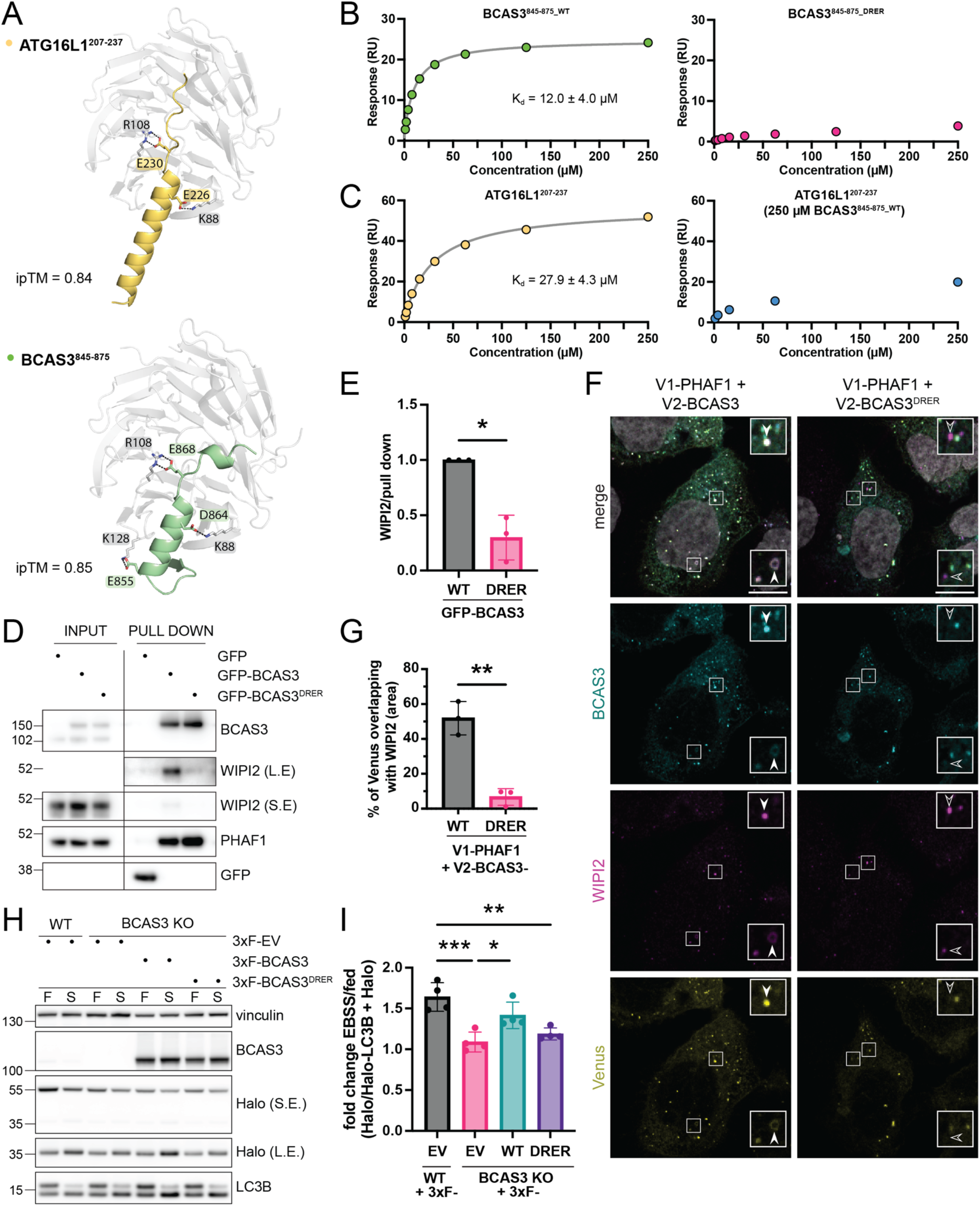
BCAS3^845-875_WT^ binds directly to WIPI2b with micromolar affinity. (**A**) AlphaFold3 models of (top) amino acids 207-237 of ATG16L1 (yellow) or (bottom) amino acids 845-875 of BCAS3 (green) with WIPI2b (grey). ipTM = 0.84 and 0.85 respectively. Residues predicted to be involved in interactions are shown as sticks. (**B**) Representative binding curve of BCAS3^845-875_WT^ (left) or BCAS3^845-875_DRER^ (right) titrated against immobilised WIPI2b^13-362Δ6CD^ using SPR. One site binding model fit is displayed as a continuous line. K_D_ = mean ± SD, n = 3. (**C**) Representative binding curve of ATG16L1^207-236^ titrated against immobilised WIPI2b^13-362Δ6CD^ in the absence (left) or presence (right) of 250 µM BCAS3^845-875_WT^ using SPR. One site binding model fit is displayed as a continuous line. K_D_ = mean ± SD, n = 3. (**D**) Representative western blot of GFP-trap. HEK293A cells were transiently transfected with either a GFP empty vector control (GFP), GFP-BCAS3 WT (GFP-BCAS3) or GFP-BCAS3^DRER^ and cultured in EBSS for 2 hours. Input samples represent 1% of the total lysate. (**E**) Quantification of (D) with pull down of WIPI2 normalised to BCAS3 pull down relative to GFP-BCAS3 WT. n = 3, Welch’s t-test, *p = 0.0266. (**F**) Representative images of HEK293A WT cells transiently transfected with V1-PHAF1 and either V2-BCAS3 WT (V2-BCAS3) or V2-BCAS3^DRER^ (Venus, yellow). Cells were treated with EBSS for 2 hours then stained with antibodies against BCAS3 (cyan) and WIPI2 (magenta). Regions of interest are shown as inset zooms. Solid arrowheads indicate co-localisation of Venus/BCAS3 and WIPI2, hollow arrowheads indicate Venus/BCAS3 signal which does not co-localise with WIPI2. Scale bar, 10 µm. (**G**) Quantification of percentage area of Venus signal which overlaps with WIPI2 signal from (F). n = 3, Welch’s t-test, **p = 0.0057. (**H**) Representative western blot of Halo-LC3B processing assay with HEK293A WT^Halo-LC3B^ or BCAS3 KO^Halo-LC3B^ cells transfected with EV, 3xFLAG-BCAS3 or 3xFLAG-BCAS3^DRER^ cultured in complete media (F) or EBSS (S) for 4 hours. (**I**) Quantification of fold change in Halo processing following starvation from (H). n = 4, one-way ANOVA followed by Tukey’s post-hoc test. ***p = 0.0006, **p = 0.0029, *p = 0.0271.

Visual comparison of these two models shows a striking similarity between their binding modes. An ATG16L1^207-236^ peptide has been shown to bind directly to WIPI2b with micromolar affinity [40]. We therefore sought to establish whether the same region of BCAS3 binds directly to WIPI2b and, if so, determine the affinity of this interaction by surface plasmon resonance (SPR). A peptide corresponding to residues 845-875 of BCAS3 (BCAS3^845-875_WT^) was added in increasing concentrations up to 250 µM (**Figure 3B**) and binding to purified WIPI2b^13-362Δ6CD^ was measured. Addition of BCAS3^845-875_WT^ resulted in a concentration-dependent response, indicating direct binding of this region of BCAS3 to WIPI2b with a K_d_ of 12.0 ± 4.0 µM. Having demonstrated that this region of BCAS3 binds directly to WIPI2b, we asked if BCAS3 residues D864 and E868 mediate its interaction with WIPI2b as indicated by the AF3 model (**Figure 3A).** Comparison of the BCAS3 protein sequence in different species showed a conservation of these residues in *Homo sapiens*, *Mus musculus*, *Danio rerio* and *Drosophila melanogaster* homologs, suggesting they might be key for BCAS3 function (**Fig. S3B**). Residues D864 and E868 in BCAS3^845-875_WT^ were substituted for arginine to generate BCAS3^845-875_DRER^ and this peptide was tested for binding to WIPI2b^13-362Δ6CD^ using SPR as before. We were unable to detect a binding response within the concentration range tested (0.98-250 µM, **Figure 3B**) suggesting that, as for ATG16L1, these residues are essential for binding of BCAS3 to WIPI2b. This is supported by a loss of BCAS3-PHAF1 binding to the WIPI2b RERE mutant (which affects the corresponding site on WIPI2b) [22].

To confirm that ATG16L1^207-237^ and BCAS3^845-875_WT^ are binding to the same site on WIPI2b, we performed a competition assay using SPR. First, we measured binding of the peptide ATG16L1^207-237^ to WIPI2b^13-362Δ6CD^ and estimated a K_d_ of 27.9 ± 4.3 µM (**Figure 3C**), which is in line with the low micromolar affinity previously reported [40]. Binding of ATG16L1^207-237^ to WIPI2b^13-362Δ6CD^ was then measured in the presence of saturating concentrations of BCAS3^845-875_WT^ (250 µM); the response was strongly decreased, indicating a significantly reduced apparent affinity. This suggests that ATG16L1^207-237^ and BCAS3^845-875_WT^ bind to the same binding site on WIPI2b (**Figure 3C**). Our data suggests that BCAS3 may bind to WIPI2b with slightly greater affinity than ATG16L1; the additional polar contact predicted to form between BCAS3^845-875^ and WIPI2b may stabilise this interaction (**Figure 3A and Fig. S3B**).

Having established D864 and E868 as key residues for binding of the BCAS3^845-875_WT^ peptide, we wanted to test whether these residues are important for interaction of the full-length proteins in cells. HEK293A cells were transfected with either a GFP empty vector control, GFP-BCAS3 WT, or the mutant GFP-BCAS3^DRER^ before 2 hours of EBSS starvation. As previously shown, GFP-BCAS3 WT co-immunoprecipitates both PHAF1 and WIPI2 (**Figure 3D**). Pull down with GFP-BCAS3^DRER^, on the other hand, showed a marked reduction in the pulldown of WIPI2 (**Figure 3D and 3E**). The interaction between GFP-BCAS3^DRER^ and PHAF1 was maintained (**Figure 3D**), suggesting that mutation of these residues does not disrupt BCAS3 tertiary structure and further confirming that the interaction between BCAS3 and PHAF1 is independent of WIPI2, as shown in **Fig. S1F**. To confirm this, V1-PHAF1 was expressed with either V2-BCAS3 or V2-BCAS3^DRER^. HEK293A WT cells were starved for 2 hours with EBSS to induce autophagy and then visualised by confocal microscopy. Both V1-PHAF1+V2-BCAS3 and V1-PHAF1+V2-BCAS3^DRER^ showed formation of Venus puncta (**Figure 3F**) confirming our observation that PHAF1 and GFP-BCAS3^DRER^ still co-immunoprecipitate (**Figure 3D**). In addition, as shown in Figure 1A, immunofluorescent staining of WIPI2 showed that approximately ∼50% of Venus-positive V1-PHAF1+V2-BCAS3 puncta colocalized with WIPI2 (**Figure 3G**). V1-PHAF1+V2-BCAS3^DRER^ showed a significant reduction in colocalization, with only around 6% of V1-PHAF1+V2-BCAS3^DRER^ Venus puncta overlapping with WIPI2 puncta (**Figure 3G**). V1-PHAF1+V2-BCAS3^DRER^ Venus puncta were generally observed to be smaller than V1-PHAF1+V2-BCAS3 which may indicate that there is a positive feedback loop which occurs when WIPI2b can interact with BCAS3.

Finally, we wanted to ascertain whether the interaction between BCAS3 and WIPI2b is functionally important for autophagy using the Halo-LC3 processing assay. HEK293A WT^Halo-LC3B^ or BCAS3 KO^Halo-LC3B^ cells were starved with EBSS for 4 hours (**Figure 3H**). BCAS3 KO^Halo-LC3B^ cells expressing a 3xFLAG empty vector showed a significant reduction in processing which could be rescued by expression of 3xFLAG-BCAS3 (**Figure 3H and 3I**). BCAS3 KO^Halo-LC3B^ cells expressing 3xFLAG-BCAS3^DRER^ showed a significant reduction in processing compared to both WT^Halo-LC3B^ and BCAS3 KO^Halo-LC3B^ rescued with 3xFLAG-BCAS3 (**Figure 3H and 3I**). The interaction between BCAS3 and WIPI2b is therefore required for starvation-induced autophagy.

Here we describe BCAS3 as a novel direct interactor of WIPI2b. The BCAS3-PHAF1 complex acts upstream of WIPI2b and, via BCAS3, can recruit it to clustered and extended structures in a PtdIns3P-dependent manner. Although WIPI2b has its own PtdIns3P binding motifs and an amphipathic helix, its membrane association seems to be additionally controlled by another factor [9,11]; we therefore propose that the BCAS3-PHAF1 complex may help to direct WIPI2b to a specific subset of membranes or stabilise it in a particular orientation at the membrane. We demonstrate that BCAS3 binds to WIPI2b in the ATG16L1-binding cleft using a similar mode of interaction but with slightly higher affinity. Our models and SPR data suggest that BCAS3 and ATG16L1 cannot bind WIPI2b at the same time; we therefore propose that there is a switch that regulates which partner WIPI2b interacts with. There are additional phosphorylation sites on WIPI2b with uncharacterised function [9]; further investigation into whether these modulate interactions with either ATG16L1 or BCAS3 will be necessary. Recently, a number of other interactors have been proposed to bind this region of WIPI2 [18,19] and there is also evidence that WIPI2 may interact with a member of the PtdIns3K complex [17] suggesting that the currently held view of WIPI2 solely to recruit ATG16L1 may be oversimplified. The function of different WIPI2-containing complexes and how the switch between them is regulated remains to be investigated.

## Methods

### Cell cultures and transfections

HEK293A cells were cultured in high-glucose DMEM (Dulbecco’s modified Eagle’s medium; Sigma, D6429) supplemented with 10% foetal bovine serum (Gibco, A52567), and penicillin-streptomycin (100 U/ml, 0.1 mg/ml respectively). Cells were grown at 37°C with 10% CO2 in a water-saturated incubator. For induction of autophagy by amino acid starvation, cells were washed and incubated in Earle’s Balanced Salt Solution (EBSS; 122 mM NaCl, 26 mM NaHCO_3_, 5.3 mM KCl, 5 mM glucose, 1 mM CaCl_2_, 0.81 mM MgSO_4_, 0.79 mM NaH_2_PO_4_) for the time indicated. Where indicated, cells were treated with 100 nM Bafilomycin A1 for 2 h.

BCAS3 KO cell lines were generated by CRISPR/Cas9. Synthetic gRNA was selected from the Thermo Fisher Scientific TrueGuide library and checked for off-target binding using the IDT guide RNA checker. The selected target sequence was in exon 6 (5’-TCGACATGGCCCAATTCGAG-3’; Thermo Fisher, A35533). HEK293A cells were transfected with gRNA according to manufacturer’s instructions using Lipofectamine^TM^ CRISPRMAX^TM^ (Invitrogen, CMAX0003) and TrueCut HiFi Cas9 protein (Invitrogen, A50575). Pools were single cell sorted by FACS using propidium iodide as a live/dead stain (Flow Cytometry STP, The Francis Crick Institute); clones were then screened by western blot and confirmed by Illumina MiSeq amplicon sequencing (Genomics STP, The Francis Crick Institute). WIPI2 KO cells were previously generated and validated [9]. ATG16L1 KO cells were kindly gifted by Anne Simonsen [16].

For transient transfection of cells, plasmid DNA (**Table 1**) was used with Lipofectamine 2000 (Invitrogen, 11668019) according to manufacturer’s instructions.

**Table 1.** DNA plasmids used for generation of cell lines.

| Name | Vector | Source |
|---|---|---|
| 3xFLAG-BCAS3 | pDEST 3xFLAG | This study |
| 3xFLAG-BCAS3<sup>R401A</sup> | pDEST 3xFLAG | This study |
| 3xFLAG-ORF | pDEST 3xFLAG | Jain et al., 2010 [42] |
| 3xFLAG-PHAF1 | pDEST 3xFLAG | This study |
| 6xHis-SUMO-3C-Twin-Strep-WIPI2<sup>13-362Δ6CD</sup> | pBR332 | This study |
| GFP-BCAS3<sup>DRER</sup> | pcDNA-DEST53 | This study |
| GFP-BCAS3<sup>WT</sup> | pcDNA-DEST53 | This study |
| GFP-EV | pEGFPC1 | Clontech |
| GFP-WIPI2a | pEGFPC1 | Dooley et al., 2014 [15] |
| GFP-WIPI2b<sup>S284D</sup> | pEGFPC1 | Gubas et al., 2024 [9] |
| GFP-WIPI2b<sup>S68D</sup> | pEGFPC1 | Gubas et al., 2024 [9] |
| GFP-WIPI2b<sup>WT</sup> | pEGFPC1 | Dooley et al., 2014 [15] |
| pMRX-IP-Halo7-LC3B | pMRX | Addgene #184899 |
| pCG-gag-pol | pCG | Yamamoto and Mizushima, 2024 [43] |
| pMD2.G-VSV.GCMV | pMD2.G | Addgene #12259 |
| V1-ORF | pcDNA3.1+ | Addgene #73635 |
| V1-PHAF1 | pcDNA3.1+ | This study |
| V1-WIPI2a | pcDNA3.1+ | This study |
| V1-WIPI2b | pcDNA3.1+ | This study |
| V2-ORF | pcDNA3.1+ | Addgene #73636 |
| V2-BCAS3<sup>DRER</sup> | pcDNA3.1+ | This study |
| V2-BCAS3<sup>WT</sup> | pcDNA3.1+ | This study |

### Plasmids

BCAS3 ORF (Origene, RC217141; corresponding to GenBank NM_001099432) and PHAF1 ORF (Origene, RG202484; corresponding to GenBank NM_025187) were used to generate 3xFLAG-BCAS3, GFP-BCAS3, 3xFLAG-PHAF1 and all WT V1- and V2-constructs by performing Gateway cloning according to manufacturer’s instructions into the corresponding pDest vector. Site-directed mutagenesis of BCAS3 was performed using the Q5 High-Fidelity PCR kit (New England Biolabs, E0554S) in the 3xFLAG-BCAS3 vector and then subcloned into other vectors using Gateway cloning. The plasmid for recombinant expression of 6xHis-SUMO-3C-Twin-Strep-WIPI2^13-362Δ6CD^ was generated by In-Fusion Snap Assembly (Takara, 638948) to assemble tags into a pBR332 expression plasmid (gift from James Le Cornu) and remove the WIPI2b 6CD loop (aa 256-297).

### Antibodies

Mouse antibodies: anti-WIPI2 ([11]), anti-FLAG M2 (Sigma, F3165), anti-vinculin (Sigma, V9264), anti-Halo (Promega, G9211), anti-GFP (CRUK, 3E1). Rabbit antibodies: anti-BCAS3 (Abcam, ab71162), anti-LC3B (Cell Signalling Technology, 2775), anti-ATG16L1 (Cell Signalling Technology, 8089S), anti-FLAG (Sigma, F7425), anti-PHAF1 (anti-c16orf70; Abcam, ab181987), anti-ULK1 (Cell Signalling Technology, 8054).

Secondary antibodies for IF: anti-rabbit IgG Alexa Fluor 488 (Invitrogen, A21206), anti-rabbit IgG Alexa Fluor 555 (Invitrogen, A31572), anti-rabbit IgG Alexa Fluor 647 (Invitrogen, A31573), anti-mouse IgG Alexa Fluor 488 (Invitrogen, A21202) and anti-mouse IgG Alexa Fluor 555 (Invitrogen, A31570). Secondary antibodies for WB: anti-mouse IgG HRP and anti-rabbit IgG HRP (Cytiva, 10196124 and 10794347 respectively).

### Western blotting

Cells were lysed with ice-cold TNTE lysis buffer (20 mM Tris-HCl pH 7.5, 150 mM NaCl, 5 mM EDTA, 1% Triton-X) containing 1X cOmplete protease inhibitor cocktail (Roche, 5056489001) and 1X PhosSTOP (Roche, 4906837001). Lysates were cleared by centrifugation, adjusted with 5X sample buffer (15% SDS [w/v], 213.5 mM Trish-HCl pH 6.8, 50% glycerol [v/v], 15% β-mercaptoethanol [v/v], 0.1% bromophenol blue [v/v]) to normalise protein concentrations and then incubated at 100°C for 10 mins. Proteins were resolved on Nu-Page Bis-Tris 4-12% gels (Life Technologies) and then transferred onto a 0.45 µM PVDF membrane (Merck). Membranes were incubated with primary and secondary antibodies and developed by enhanced chemiluminescence (Classico or Crescendo Western HRP substrate [Merck Millipore, WBLUC0100 and WBLUR0100 respectively]); for antibodies with weak signal, blots were developed using ECL Select^TM^ Western Blotting Detection Reagent (Cytiva, GERPN2235). Visualisation of antibody signal was performed using an Amersham ImageQuant 800 imager (Cytiva). Quantification of western blot signal was performed using densitometry with FIJI [44] or Image Lab (BioRad).

### Halo-LC3B processing assay

Cell lines stably expressing Halo-LC3B were generated according to previously published protocols [43,45] with minor changes. For retrovirus generation, HEK293FT cells were transfected with pMRX-IP-Halo7-LC3B (gift from Noboru Mizushima), pCG-gag-pol, and pMD2.G-VSV.GCMV (gift from Didier Trono) using Lipofectamine 2000 (Invitrogen, 11668019). After two days, virus containing medium was harvested, filtered using a 0.45 µm filter and aliquots were frozen at -80°C. To generate cell lines stably expressing Halo-LC3B, HEK293A WT and BCAS3 KO cells were transduced with Halo-LC3B retroviral supernatant with 8 µg/ml Polybrene (Merck, TR-1003-G). The following day, selection medium containing 0.5 µg/ml puromycin (Gibco, A1113803) was added. Expanded cells were stained with Halo-TMR ligand (Promega, G8252) and sorted by FACS (Flow Cytometry Science Technology Platform, The Francis Crick Institute) to obtain a pool with a consistent expression level. For Halo-LC3B processing assays, cells were treated with 10 µM of the Halo ligand 7-Bromo-1-Heptanol (Thermo Fisher Scientific, H54762.03) for 20 minutes. Cells were washed thrice and incubated with EBSS or DMEM for the time indicated. Halo-LC3B processing was assessed by western blot; densitometry was used to quantify free Halo signal and Halo-LC3B signal with Halo processing being expressed as a ratio of free Halo:(free Halo + Halo-LC3B).

### Immunoprecipitation/GFP-trap

Cells were lysed as for western blot. Cleared lysates were then incubated on agarose binding control beads (ChromoTek, bab-20) for 1 h, rotating at 4°C. Resulting supernatant was then further incubated on GFP-trap beads (ChromoTek, gta-20) for 2 h, rotating at 4°C. Bound samples were washed in TNTE wash buffer (20 mM Tris-HCl pH 7.5, 150 mM NaCl, 5 mM EDTA, 0.1% Triton-X) and proteins were eluted with 2X Laemmli sample buffer at 65°C for 5 mins before resolving by SDS-PAGE and western blotting.

### Immunofluorescence

Cells were seeded on #1.5 glass coverslips. Fixation was performed with 3% PFA (16% PFA [CliniSciences, 15710-S] diluted with 0.1 mM CaCl_2_, 0.1 mM MgCl_2_ in PBS-A) for 20 mins at room temperature. Fixation was quenched by washing and incubating with 50 mM ammonium chloride for 10 mins. For cells to be stained with antibodies, cells were permeabilized with 0.2% Triton-X (Fisher Scientific, 11488696) for 3-5 mins at room temperature. Following permeabilization, coverslips were blocked in 1% BSA (Sigma-Aldrich, A9647) for 30 minutes. Primary antibody (in 1% BSA) incubation was performed for 1 h, followed by three washes in 1% BSA and then secondary antibody (in 1% BSA) incubation for 1 h. Nuclear staining with Hoechst 33342 (5 µg/ml in PBS-A [137 mM NaCl, 10 mM Na_2_HPO_4_, 3.35 mM KCl, 1.8 mM KH_2_PO_4_]; Invitrogen, H3570) was performed for 5 mins before final washes in PBS and water. Coverslips were mounted onto SuperFrost slides (Thermo Scientific) with MOWIOL 4-88 (Millipore, 475904-M).

### Imaging and analysis

An LSM 880 confocal microscope (Zeiss) fitted with a 63x/1.4 NA oil immersion objective was used to image coverslips. Image analysis was performed using semi-automated pipelines. A FIJI [44] macro was used to batch pre-process raw data ready for analysis. In brief, an Ilastik [46] model was trained to detect relevant puncta or structures (i.e WIPI2 puncta, WIPI2 structures, BCAS3/PHAF1 (3xFLAG) structures or Venus puncta as indicated) and binary masks of segmented features were produced using FIJI. A CellProfiler 4 [47] pipeline was then used to segment nuclei and cells (either all cells within a field or only transfected cells where relevant) and calculate either the density of puncta/structures within the cytoplasmic area (“nucleus” subtracted from “cell”) or the overlap between two segmented masks (i.e. percentage of Venus-positive area overlapping with WIPI2-positive area).

### Expression and Purification of Twin-Strep-WIPI2b^13-362Δ6CD^

Expression and purification of Twin-Strep-WIPI2b^13-362Δ6CD^ was adapted from Gong et al., 2023 [40]. 6xHis-SUMO-3C-Twin-Strep-WIPI2^13-362Δ6CD^ was expressed in *E. coli* Rosetta 2(DE3) pLysS (Merck, 71403-3) in TB medium after induction with 100 µM IPTG (Melford, I56000-25.0) for 22 h at 18°C. Subsequently, bacteria were harvested in lysis buffer (50 mM Na_2_HPO_4_/NaH_2_PO_4_ pH 7.5, 300 mM NaCl, 5% glycerol, 10 mM Imidazole, 0.5% Triton X-100, 0.5 mM TCEP, 25 U/ml Benzonase, 2 mM MgCl_2_, 1X cOmplete EDTA-free Protease Inhibitor [Roche, 5056489001]) and lysed using a Cell Disruptor (French Press). After the lysate was cleared by centrifugation (45 mins, 45,000g), the supernatant was incubated with HisPur Cobalt Resin (Thermo Fisher Scientific, 89965) for 1.5 h at 4°C. The resin was collected in an empty column, washed with wash buffer (50 mM Na_2_HPO_4_/NaH_2_PO_4_ pH 7.5, 300 mM NaCl, 5% Glycerol, 0.5 mM TCEP, 1X cOmplete EDTA-free Protease Inhibitor [Roche, 5056489001]) and subsequently eluted with increasing concentrations of imidazole (50, 75, 100 & 150 mM) in the wash buffer. The four fractions were pooled, digested with GST-His-3C protease (Structural Biology Science Technology Platform, The Francis Crick Institute), while dialyzing to the size exclusion chromatography (SEC) buffer (25 mM Tris HCl pH 7.5, 150 mM NaCl, 0.5 mM TCEP, 3% Glycerol, 1 mM MgCl_2_) using a 3,500 MWCO Slide-A-Lyzer cassette (Thermo Fisher Scientific, 66110) overnight at 4°C. The GST-His-3C protease was removed by incubation with GSH-Sepharose (Cytiva, 17075605) for 1h at 4°C, which was then removed by centrifugation and filtration. The sample was concentrated using a 10,000 MWCO Amicon Ultra Centrifugal filter (Merck Millipore, UFC901024) and further purified by SEC using a Superdex 200 16/600 column (GE Healthcare). Fractions containing Twin-Strep-WIPI2b^13-362Δ6CD^ were pooled, incubated with HisPur Cobalt Resin to remove remaining His-Sumo and concentrated, before being snap frozen in aliquots in liquid N_2_ and stored at - 80°C.

### Surface plasmon resonance

Surface plasmon resonance (SPR) was used to determine the affinity between purified Twin-Strep-WIPI2b^13-362Δ6CD^ and 31 amino acid peptides, ATG16L1^207-237^ and BCAS3^845-875^, spanning the predicted WIPI2b binding sites. Peptides were synthesized using FMOC solid-phase peptide chemistry by Dhira Joshi (Chemical Biology Science Technology Platform, The Francis Crick Institute) and can be found in **Table 2**. Experiments were carried out on a Biacore S200 (Cytiva) at 25°C in PBS-P+ buffer (20 mM phosphate pH 7.4, 2.7 mM KCl, 137 mM NaCl, 0.05% Tween 20, 1 mM MgCl_2_, and 0.5 mM TCEP). Strep-Tactin (Twin-Strep-tag Capture Kit; IBA Lifesciences, 2-4370-000) was immobilised on a Series S Sensor Chip CM5 (Cytiva, 29149604) using amine coupling chemistry. Twin-Strep-WIPI2b^13-362Δ6CD^ (10-50 nM) was then captured and peptides dilution series (0.98, 1.95, 3.91, 7.81, 15.63, 31.25, 62.5, 125, 250 µM) were injected on to the Twin-Strep-WIPI2b^13-362Δ6CD^ loaded flow channel as well as onto a reference channel. Flow rate was 30 μl/min and association and dissociation were recorded. For the competition assay, saturating concentrations of BCAS3^845-875^ (250 µM) were injected, followed by samples containing ATG16L1^207-237^ at increasing concentrations (0.98, 3.91, 15.63, 62.50, 250 µM) and 250 µM BCAS3^845-875^. Reference subtracted data was analysed by plotting the instrument response over peptide concentration and fitting data using the steady state affinity (1:1) model in the Biacore S200 Evaluation software to estimate equilibrium dissociation constants (K_d_).

**Table 2.** Synthetic peptide sequences generated for use in surface plasmon resonance experiments.

| **Peptide Name** | **Sequence** |
|---|---|
| ATG16L1<sup>207-237</sup> | NH<sub>2</sub>- AENEKDSRRRQARLQKELAEAAKEPLPVEQD-CONH<sub>2</sub> |
| BCAS3<sup>845-875_WT</sup> | NH<sub>2</sub>-LRHMSSMEHTEEGLRERLADAMAESPSRDVV-CONH<sub>2</sub> |
| BCAS3<sup>845-875_DRER</sup> | NH<sub>2</sub>-LRHMSSMEHTEEGLRERLA**R**AMA**R**SPSRDVV-CONH<sub>2</sub> |

### AlphaFold3 modelling

AlphaFold 3 [38] was used for structural prediction of the BCAS3-PHAF1-WIPI2b complex as well as ATG16L1^207-237^-/BCAS3^845-875^-WIPI2b complexes using the AlphaFold Server with default settings. For WIPI2b, BCAS3 and PHAF1, full-length sequences as provided on UniProt were used as input (UniProt: Q9Y4P8, Q9H6U6, and Q9BSU1 respectively). PAE, iPTM and PTM values were inspected manually and the model with the highest ranking score was analysed and visualised using PyMOL 3.1.6.1.

### Statistical analysis

Statistics were performed using GraphPad Prism 11 software as detailed in the figure legends. Unless otherwise specified, data are presented as mean ± SD.

## Supplementary Figure Legends

**Figure S1.**
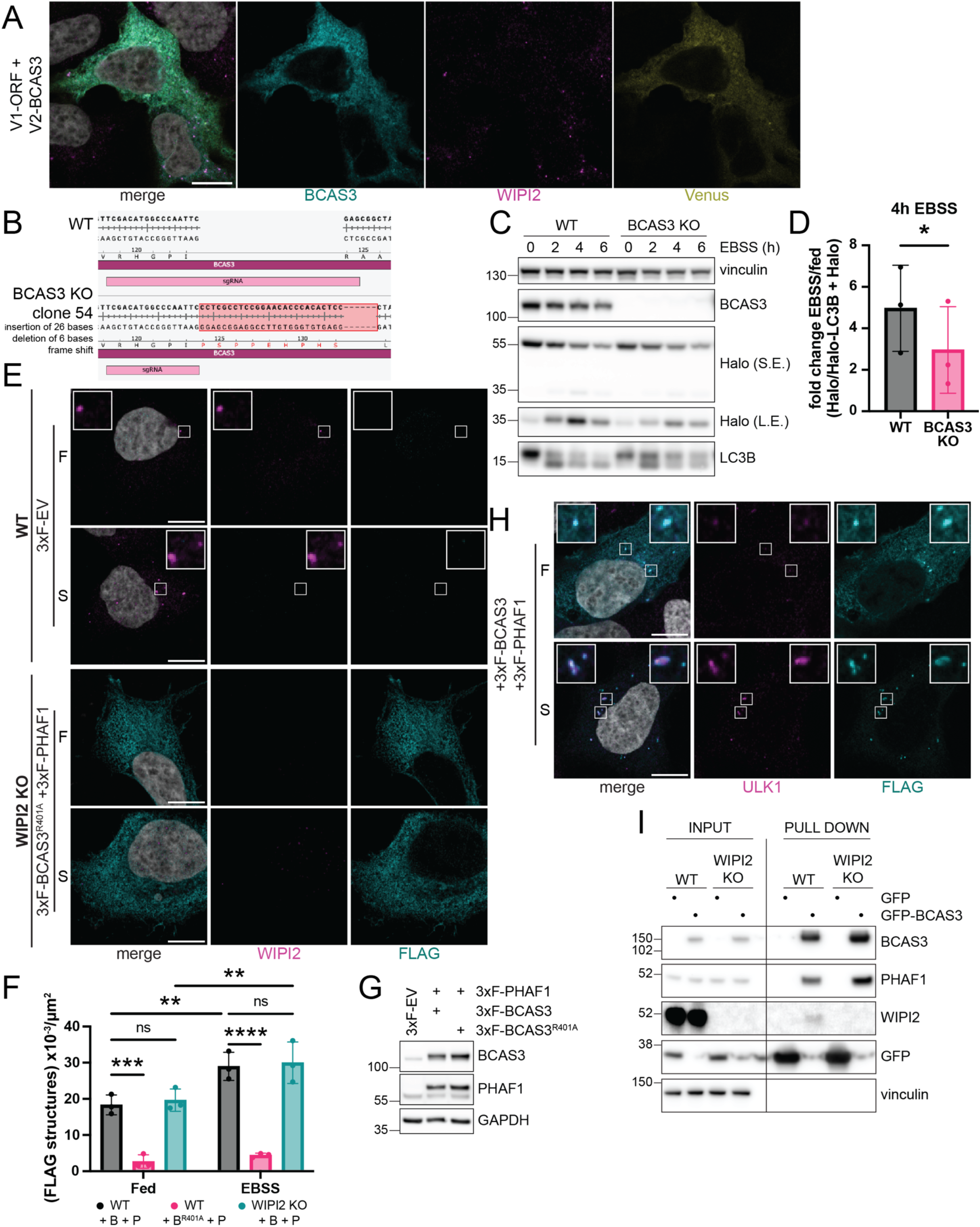
Generation and characterisation of HEK293A BCAS3 KO cells. (**A**) Representative images of HEK293A WT cells transiently transfected with V1-ORF and V2-BCAS3 for split Venus assay (yellow). Cells were treated with EBSS for 2 hours then stained with antibodies against BCAS3 (cyan) and WIPI2 (magenta). Scale bar, 10 µm. (**B**) Amplicon sequencing of HEK293A WT and BCAS3 KO clone 54. BCAS3 KO clone 54 shows an insertion of 26 bases and a deletion of 6 bases resulting in a frame shift and insertion of a stop codon in exon 6. (**C**) Representative western blot of Halo-LC3B processing assay with HEK293A WT^Halo-LC3B^ or BCAS3 KO^Halo-LC3B^ cells cultured in complete medium or EBSS for 0, 2, 4 or 6 hours as indicated. (**D**) Quantification of fold change in Halo processing following 4 hours starvation from (C). n = 3, paired t-test. *p = 0.0459. (**E**) Representative images of (top) HEK293A WT cells transiently transfected with a 3xFLAG empty vector control (3xF-EV) or (bottom) WIPI2 KO cells transiently transfected with 3xFLAG-PHAF1 (3xF-PHAF1) and 3xFLAG-BCAS3^R401A^ (3xF-BCAS3^R401A^). Cells were cultured in complete media (F) or EBSS (S) for 2 hours and then stained with antibodies against WIPI2 (magenta) and FLAG (cyan). Regions of interest are shown as inset zooms. Scale bar, 10 µm. (**F**) Quantification of density of FLAG-positive structures as seen in Figure 1F. HEK293A WT or WIPI2 KO cells were transiently transfected with 3xFLAG empty vector, 3xFLAG-BCAS3 WT + 3xFLAG-PHAF1 (B + P), or 3xFLAG-BCAS3^DRER^ + 3xFLAG-PHAF1 (B^R401A^ + P). n = 3, two-way ANOVA followed by Tukey’s post-hoc test. [WT + B + P fed] vs [WIPI2 KO + B + P fed] ns = 0.8843, [WT + B + P EBSS] vs [WIPI2 KO + B + P EBSS] ns = 0.9307, [WT + B + P fed] vs [WT + B + P EBSS] **p = 0.0023, [WIPI2 KO + B + P fed] vs [ WIPI2 KO + B + P EBSS] **p = 0.0028, ***p = 0.0003, ****p <0.0001. (**G**) Western blot of HEK293A WT cells transiently transfected with a 3xFLAG empty vector control (3xF-EV) or 3xFLAG-PHAF1 (3xF-PHAF1) with either 3xFLAG-BCAS3 WT (3xF-BCAS3) or 3xFLAG-BCAS3^R401A^ (3xF-BCAS3^R401A^) as indicated demonstrating comparable expression levels of mutant. (**H**) Representative images of HEK293A WT cells transiently transfected with 3xFLAG-PHAF1 (3xF-PHAF1) and 3xFLAG-BCAS3 WT (3xF-BCAS3). Cells were cultured in complete media (F) or EBSS (S) for 2 hours and then stained with antibodies against ULK1 (magenta) and FLAG (cyan). Regions of interest are shown as inset zooms. (**I**) Representative western blot of GFP-trap. HEK293A WT or WIPI2 KO cells were transiently transfected with either a GFP empty vector control (GFP) or GFP-BCAS3 WT (GFP-BCAS3) and cultured in EBSS for 2 hours. Input samples represent 1% of the total lysate.

**Figure S2.**
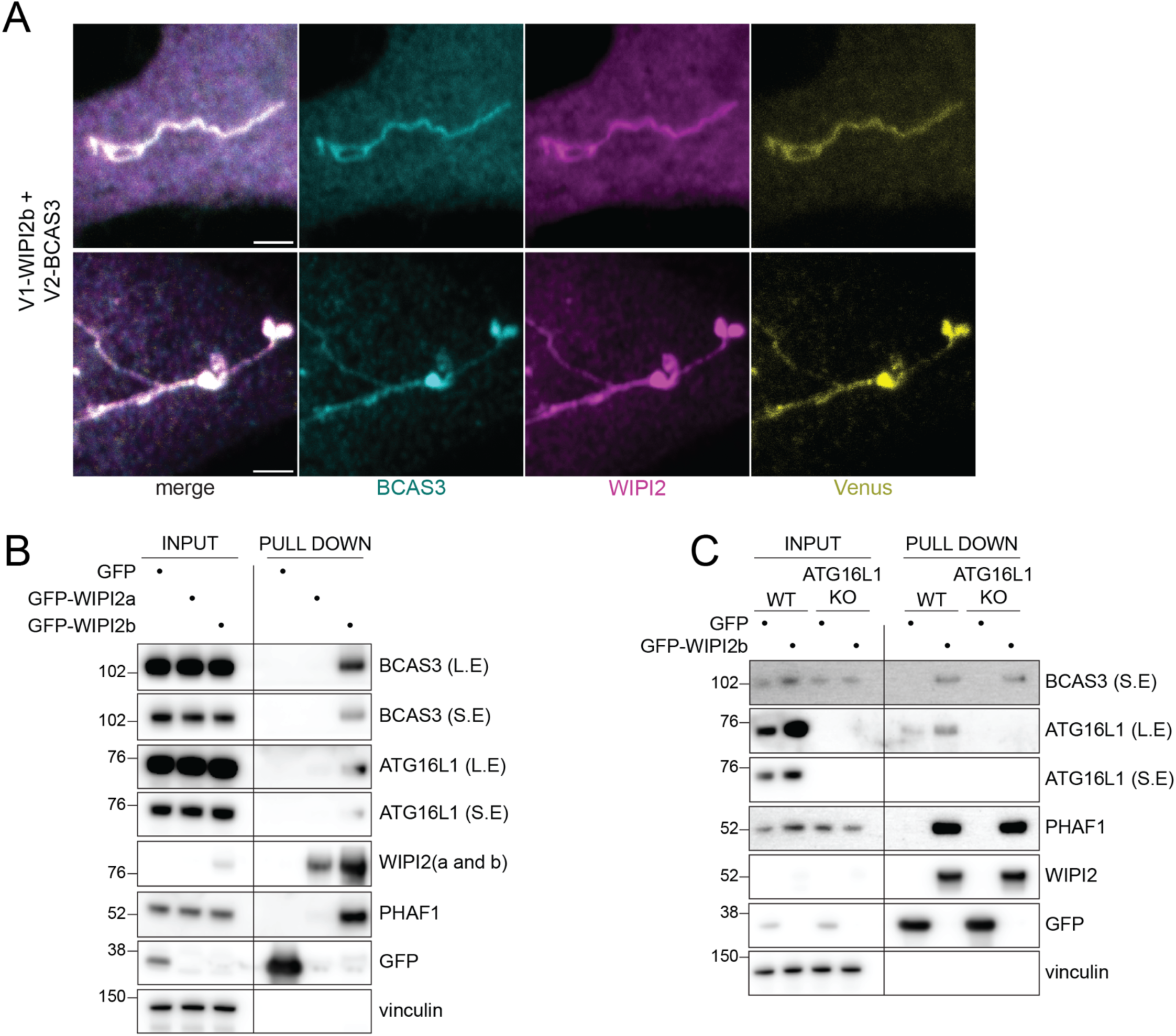
BCAS3 interacts with WIPI2b in an ATG16L1-independent manner. (**A**) Examples of particularly large extended structures observed with expression of V1-WIPI2b and V2-BCAS3 following 2 hours EBSS starvation as described in Figure 2B. Maximum projections of Z-stacks. Scale bar, 2 µm. Note: brightness and contrast adjusted separately for example images to allow visualisation. (**B**) Western blot of GFP-trap. HEK293A WIPI2 KO cells were transiently transfected with either a GFP empty vector control (GFP), GFP-WIPI2b or GFP-WIPI2a and cultured in EBSS for 2 hours. Input samples represent 1% of the total lysate. (**C**) Western blot of GFP-trap. HEK293A WT or ATG16L1 KO cells were transiently transfected with either a GFP empty vector control (GFP) or GFP-WIPI2b and cultured in EBSS for 2 hours. Input samples represent 1% of the total lysate.

**Figure S3.**
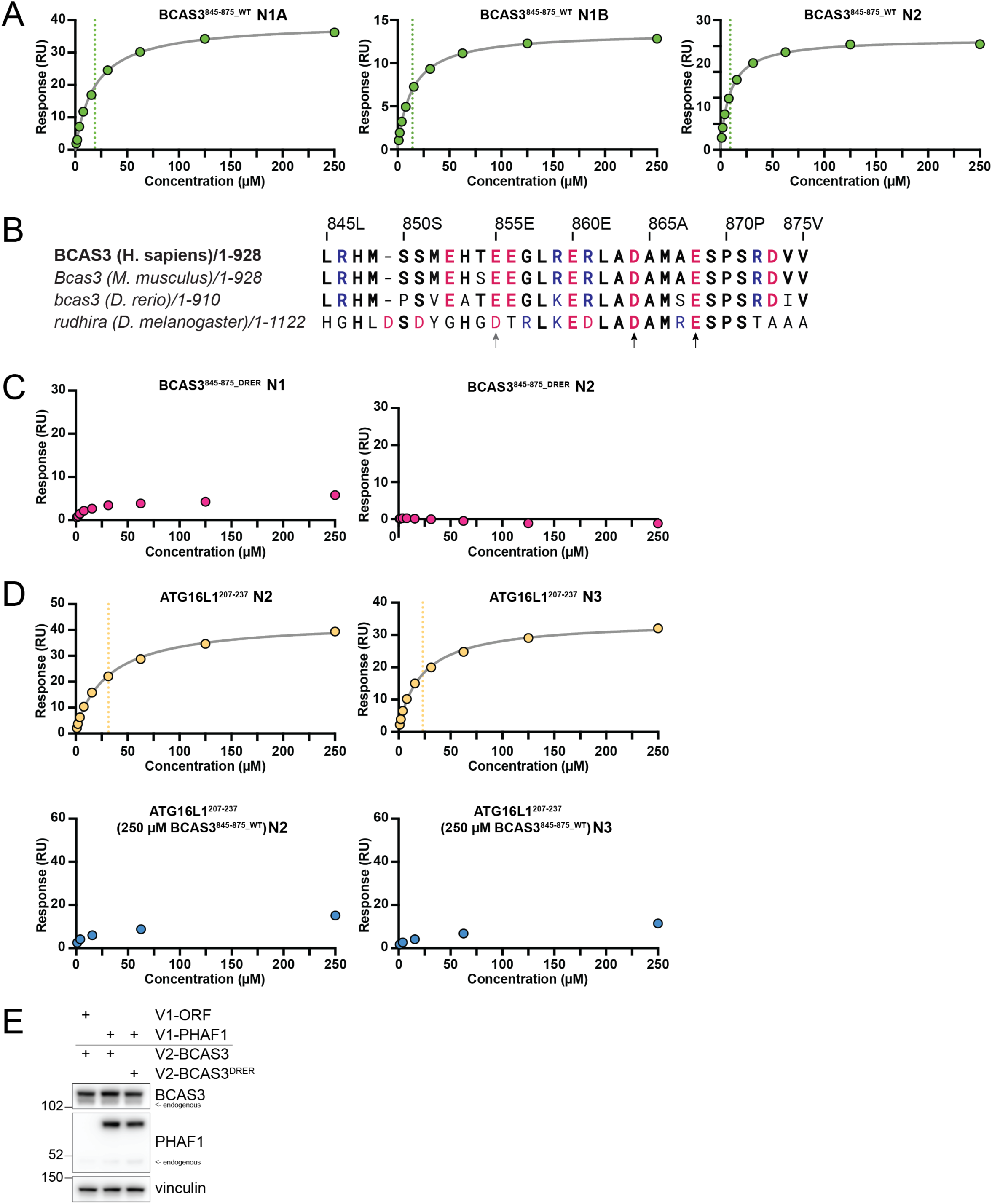
BCAS3 residues D864 and E868 are required for WIPI2b interaction (**A**) Binding curves from replicates of SPR experiments shown in Figure 3B. BCAS3^845-875_WT^ titrated against immobilised WIPI2b^13-362Δ6CD^. (**B**) Alignment of sequences of BCAS3 homologs in different species covering the BCAS3 peptide region [48]. Positively and negatively charged residues are highlighted in blue and red respectively. Bold residues indicate conservation with the *H. sapiens* homolog. Black arrows indicate residues corresponding to *H. sapiens* BCAS3 D864 and E868. Grey arrow indicates other potential interacting residue E855 based on Figure 3A. (**C**) Binding curves from replicates of SPR experiments shown in Figure 3B. BCAS3^845-875_DRER^ titrated against immobilised WIPI2b^13-362Δ6CD^. (**D**) Binding curves from different replicates of SPR experiments shown in Figure 3C. ATG16L1^207-237^ titrated against immobilised WIPI2b^13-362Δ6CD^ in the absence (top) or presence (bottom) of 250 µM BCAS3^845-875_WT^. (**A, C and D**). Where appropriate, one site binding model fit is displayed as a continuous line. Dotted line indicates K_d_ for replicate. (**E**) Western blot of HEK293A WT cells transiently transfected with V1-ORF or V1-PHAF1 and V2-BCAS3 or V2-BCAS3^DRER^ as indicated demonstrating comparable expression levels of mutant.

## Abbreviations

AF3: AlphaFold 3
BCAS3: BCAS3 microtubule associated cell migration factor
EBSS: Earle’s balanced salt solution
KO: knockout
PHAF1: phagosome assembly factor 1
PROPPIN: β-propellers that bind phosphoinositides
PtdIns3K: phosphatidylinositol 3-kinase complex I
PtdIns3P: phosphatidylinositol 3-phosphate
SPR: surface plasmon resonance
WIPI: WD repeat domain, phosphoinositide interacting
WT: wild type

